# Structural basis for a dual-function type II-B CRISPR-Cas9

**DOI:** 10.1101/2024.10.22.619592

**Authors:** Grace N. Hibshman, David W. Taylor

## Abstract

Cas9 from *Streptococcus pyogenes* (SpCas9) revolutionized genome editing by enabling programmable DNA cleavage guided by an RNA. However, SpCas9 tolerates mismatches in the DNA-RNA duplex, which can lead to deleterious off-target editing. Here, we reveal that Cas9 from *Francisella novicida* (FnCas9) possesses a unique structural feature—the REC3 clamp—that underlies its intrinsic high-fidelity DNA targeting. Through kinetic and structural analyses, we show that the REC3 clamp forms critical contacts with the PAM-distal region of the R-loop, thereby imposing a novel checkpoint during enzyme activation. Notably, *F. novicida* encodes a non-canonical small CRISPR-associated RNA (scaRNA) that enables FnCas9 to repress an endogenous bacterial lipoprotein gene, subverting host immune detection. Structures of FnCas9 with scaRNA illustrate how partial R-loop complementarity hinders REC3 clamp docking and prevents cleavage in favor of transcriptional repression. The REC3 clamp is conserved in type II-B CRISPR-Cas9 systems, pointing to a potential path for engineering precise genome editors or developing novel antibacterial strategies. These findings reveal the dual mechanisms of high specificity and virulence by FnCas9, with broad implications for biotechnology and therapeutic development.

**Graphical Abstract:** 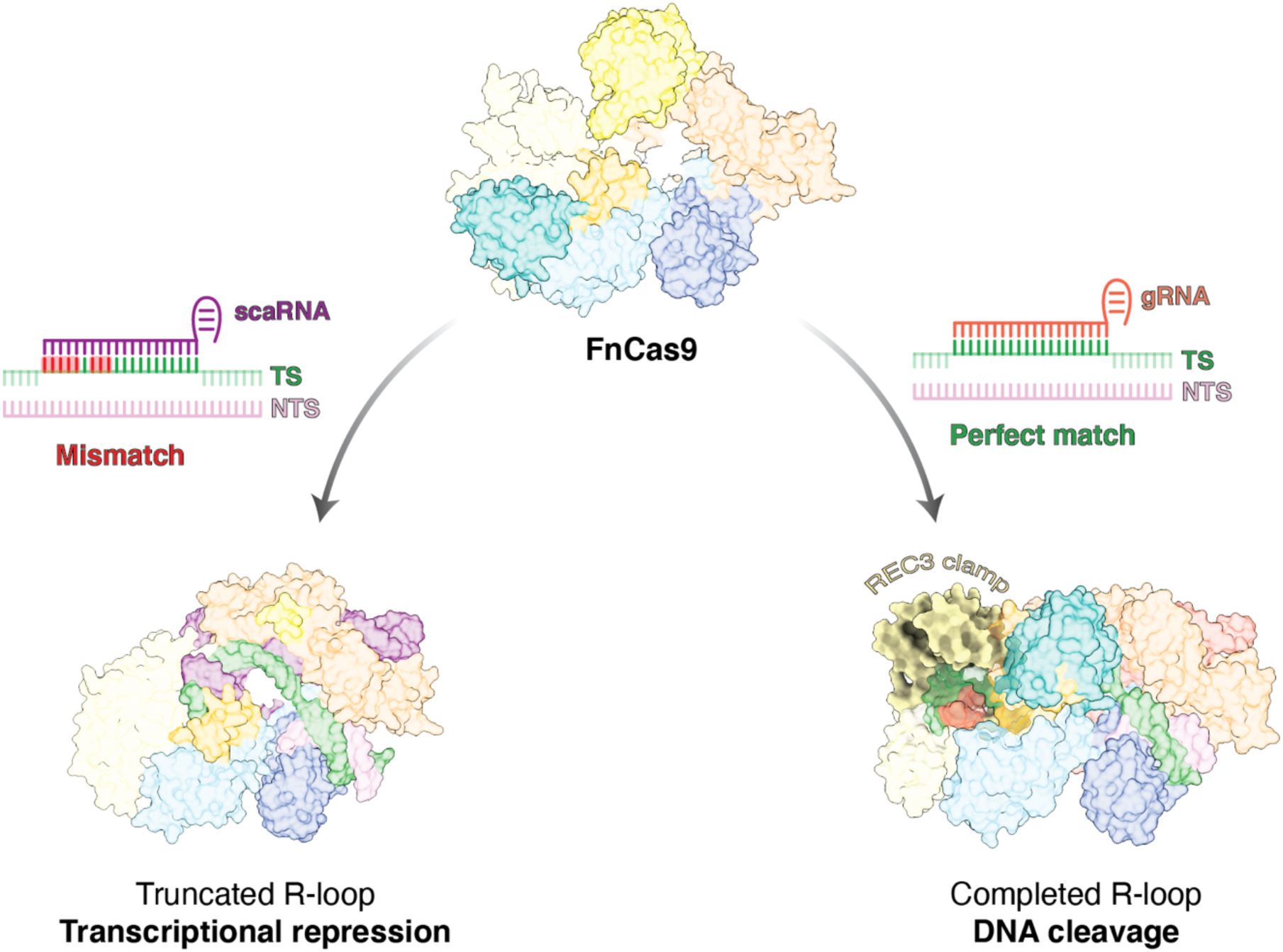

## Introduction

Bacterial and archaeal species have evolved CRISPR-Cas systems as an adaptive immune defense against foreign genetic elements, including bacteriophages and plasmids^1^. Among these systems, the Cas9 endonuclease quickly gained attention for its widespread applications in genome engineering^2–4^. *Streptococcus pyogenes* Cas9 (SpCas9) is the most used Cas9 ortholog, but suffers from off-target effects, which pose the risk of introducing unintended genomic modifications^5,6^. Multiple groups have attempted to improve SpCas9 fidelity through directed evolution and rational engineering, often trading off cleavage efficiency for reduced off-target activity ^7–11^.

SpCas9 has been repurposed for widespread applications beyond genome editing, including targeted transcriptional regulation^12–14^. To accomplish transcriptional repression, the catalytic residues of SpCas9 are mutated to produce a catalytically “dead” Cas9 (dCas9) that binds DNA without the possibility of cleavage. By fusing dCas9 to transcriptional effectors, one can direct the complex to promoter or enhancer regions for gene silencing or activation, respectively^15,16^. This approach ensures that the cleavage activity is eliminated by preventing any unintentional genomic damage.

In the search for naturally occurring high-fidelity alternatives, *Francisella novicida* Cas9 (FnCas9) emerged as an enzyme with intrinsic specificity^17^. Unlike SpCas9, which tolerates mismatches at the distal end of the RNA-DNA heteroduplex^18,19^, FnCas9 shows dramatically reduced cleavage in the presence of these same mismatches. Although early studies demonstrated this enhanced specificity, the molecular basis remains poorly understood.

Beyond its role in genome editing, *F. novicida* uniquely encodes a small CRISPR-associated RNA (scaRNA) that directs FnCas9 to repress, rather than cleave, an endogenous bacterial lipoprotein gene^20,21^. This transcriptional silencing mechanism aids *F. novicida* in evading host immune detection. Crucially, FnCas9 achieves transcriptional repression while retaining cleavage activity. FnCas9 can have dual function as both a DNA endonuclease and a transcriptional repressor in its native, catalytically active form. This is in stark contrast to SpCas9, which requires catalytic inactivation to safely mediate transcriptional control without inadvertently cleaving target DNA.

The dual functionality of FnCas9 hints at a unique mechanistic checkpoint enabling the enzyme to discriminate between targets destined for cleavage and those targeted for transcriptional repression. Here, we combine kinetic measurements of mismatch discrimination with cryo-EM structural analysis to capture distinct FnCas9 conformational states. From these data, we identify a previously unrecognized structural feature that serves as a checkpoint in FnCas9 activation that we call the REC3 clamp. We demonstrate that nearly full R-loop complementarity is required for the REC3 clamp to dock and trigger DNA cleavage. In contrast, incomplete complementarity prevents REC3 clamp docking and thereby represses transcription as visualized by cryo-EM structures of FnCas9 with scaRNA-bound targets. Thus, our findings reveal how FnCas9 accomplishes high-fidelity DNA cleavage and selective transcriptional repression.

Finally, we establish that the REC3 clamp is highly conserved across type II-B CRISPR-Cas9 enzymes, suggesting that this dual function may be a hallmark of related Cas9 orthologs. By detailing the structural and mechanistic underpinnings of FnCas9 specificity and dual functionality, our work provides a blueprint for developing high-fidelity genome editors and leveraging the scaRNA-based repression mechanism for antibacterial strategies.

## Materials and Methods

### Protein expression and purification

FnCas9 was expressed from pET-His6-FnCas9GFP purchased from Addgene (Addgene plasmid #130966; http://n2t.net/addgene:130966; RRID:Addgene_130966)^17^. FnCas9 was then expressed and purified as previously described^17^, except NiCo21(DE3) (New England Biolabs) were used instead of Rosetta2 (DE3). Additionally, after affinity purification with Ni-NTA beads, the lysate was exposed to TEV protease to liberate GFP. The remainder of the purification was executed as previously described.

FnCas9 Y794A mutant was cloned using Q5 Hot Start Hi-Fidelity polymerase and KLD kit (New England Biolabs). The mutated sequences were verified by Plasmidsaurus and purified in the same manner as WT FnCas9.

### Nucleic acid preparation

DNA duplexes 55 nt long were prepared from PAGE-purified oligonucleotides synthesized by Integrated DNA Technologies as previously described^39^. The sequences of the synthesized oligonucleotides, including the positions of mismatches, are listed in **Extended Data Table 2**.

### Buffer composition and kinetic reactions

Cleavage reactions were performed in 1X cleavage buffer (20 mM Tris-Cl, pH 7.5, 100 mM KCl, 10mM MgCl_2_, 5% glycerol, 1 mM DTT) at 37 °C.

### DNA cleavage kinetics

The reaction of FnCas9 with on- and off-target DNA was performed by preincubating FnCas9.gRNA (500 nM active-site concentration of FnCas9, 750 nM gRNA) in 5X cleavage buffer supplemented with 0.2 mg/mL molecular biology grade BSA for 10 minutes at room temperature. The reaction was initiated by the addition of 5’-FAM-labeled DNA duplex at the final concentrations of 10 nM DNA, and 100 nM preincubated enzyme.

The reactions were carried out at 37°C and stopped at various times by mixing with 0.5 M EDTA. Reaction products were resolved and quantified using an Applied Biosystems DNA sequencer (ABI 3130xl) equipped with a 36 cm capillary array and nanoPOP7 polymer (MCLab)^40^. Data fit to equations were fit using either a single or double-exponential equations shown below:

Single exponential equation:

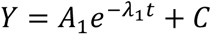

where Y represents concentration of cleavage product, A_1_ represents the amplitude, λ_1_ represents the observed decay rate (eigenvalue) and C is the endpoint. The half-life was calculated as t_1/2_ = ln(2)/λ_1_.

Double exponential equation:

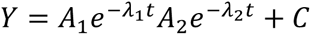

where Y represents concentration of cleavage product, A_1_ represents the amplitude and λ_1_ represents the observed rate for the first phase. A_2_ represents the amplitude and λ_2_ represents the observed rate for the second phase, and C is the endpoint.

### Cryo-EM sample preparation, data collection, and processing

FnCas9 bound to different DNA substrates were assembled by mixing FnCas9 with gRNA in a 1:1.5 molar ratio and incubated at room temperature for 10 minutes in reaction buffer (20 mM Tris-Cl, pH 7.5, 100 mM KCl, 10 mM MgCl2, 5% glycerol, and 5 mM DTT). Each DNA substrate was then added in a 1:1 molar ratio with either 8 µM FnCas9 gRNP for the on-target DNA substrate, or 10µM FnCas9 gRNP for the off-target DNA substrate and scaRNA 1101 DNA substrate. The on-target complex was incubated for 1 hour at room temperature, and the off-target complex was incubated for 2 hours at room temperature. The reactions were quenched by vitrification. 2.5 µl of sample was applied to glow discharged holey carbon grids (Quantifoil 1.2/1.3), blotted for 6 s with a blot force of 0, and rapidly plunged into liquid nitrogen-cooled ethane using an FEI Vitrobot MarkIV.

FnCas9 perfect match and mismatch datasets were collected on an FEI Titan Krios cryo-electron microscope equipped with a K3 Summit direct electron detector (Gatan, Pleasanton, CA). Images were recorded with SerialEM v4.1^41^ with a pixel size of 0.83 Å. Movies were recorded at 13.3 electrons/pixel/second for 6 s (80 frames) to give a total dose of 80 electrons/pixel. FnCas9 scaRNA 1101 DNA dataset was collected on an FEI Glacios cryo-TEM equipped with a Falcon 4 detector with a pixel size of 0.933Å, and a total exposure time of 15 s resulting in a total accumulated dose of 49 e/ Å^2^. All datasets were collected with a defocus range of −1.5 to −2.5 µm. Motion correction, CTF estimation and particle picking was performed on-the-fly using cryoSPARC Live v4.0.0-privatebeta.2^42^. Further data processing was performed with cryoSPARC v.3.2. A total of 5,378 movies were collected for the on-target dataset, 7,158 movies were collected for the off-target dataset, and 2,583 movies were collected for the scaRNA dataset.

The initial stages of processing were similar for all datasets, where blob picker was used with a minimum particle diameter of 120 Å and a maximum particle diameter of 220 Å. The off-target dataset particles were then subjected to a single round of 2D classification. The particles selected from 2D classification were processed via ab initio reconstruction, followed by heterogeneous refinement. The on-target and scaRNA dataset particles were not subjected to 2D classification, but rather, went straight into ab initio reconstruction, followed by heterogeneous refinement. The best class from heterogeneous refinement was then fed to non-uniform refinement. The mask from the non-uniform refinement was used for 3D variability, which was displayed using the cluster output mode, with 10 clusters, and a 5 Å filter resolution. The clusters were grouped by the presence of the HNH, RuvC, and REC3 domains for the on-target dataset. The particles corresponding to a single state were extracted with a 400-pixel box size for the on-target particles, 420-pixel box size for the scaRNA particles, and 384-pixel box size for the off-target particles. The extracted particles were then globally and locally CTF refined, and fed to non-uniform refinement which yielded the final reconstructions.

### Structural model building and refinement

Inactive FnCas9 (Protein Data Bank (PDB) code: 5B2O) was used as a starting model for the on-target FnCas9 structure with RuvC and HNH present. The HNH domain from the starting model was removed and replaced through rigid body fitting with a model of the HNH domain folded by AlphaFold2^43^. HNH L1 and L2 were built, and nucleic acid alterations were made in Coot v1.1.07^44^. Further modeling was performed using Isolde v1.6^45^. The model was ultimately subjected to real-space refinement implemented in Phenix v1.21^46^. The remaining on-target structures used the FnCas9 structure with RuvC and HNH present as a starting model. The productive off-target structure also used the FnCas9 structure with RuvC and HNH present as a starting model, but the non-productive off-target structure used the non-productive structure of FnCas9 with RuvC present as its starting model. The Cas9 sequence from *F. novicida U112* was supplied to the AlphaFold3 web server^29^, along with a gRNA mimicking scaRNA and a 72 bp DNA comprising the 5’UTR of *FTN_1101*. This predicted model was used as a starting model for the scaRNA gRNA 1101 DNA structure. These structures were then subjected to a similar refinement workflow, where nucleic acid alterations were made in Coot v1.1.07^44^, modeling was done in Isolde v1.6^45^, and ultimately subjected to real-space refinement implemented in Phenix v1.21^46^. All structural figures were generated using ChimeraX v1.2^47^.

### Fluorescence anisotropy binding assays

FnCas9 gRNP complexes were prepared as described above, and serial 2-fold dilutions were incubated with 0.8 nM FAM-labeled phosphorothioate DNA for 2h at 37 °C in 1 x cleavage buffer supplemented with 0.05% Tween-20. Data were recorded at 37 °C in a CLARIOstar Plus multi-detection plate reader (BMG Labtech) equipped with a fluorescence polarization optical module (λ_ex_ = 485 nm; λ_em_ = 520 nm). The data were fit using a one-step binding model in KinTek Explorer to define the K_d_ and the extrapolated starting (Y_0_) and ending (ΔY) points, which were used to normalize the data for calculation of fraction bound (**Fig. 2f**):

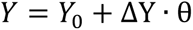

and

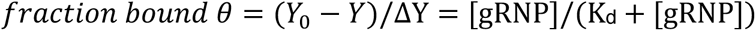

**Fig. 1.**
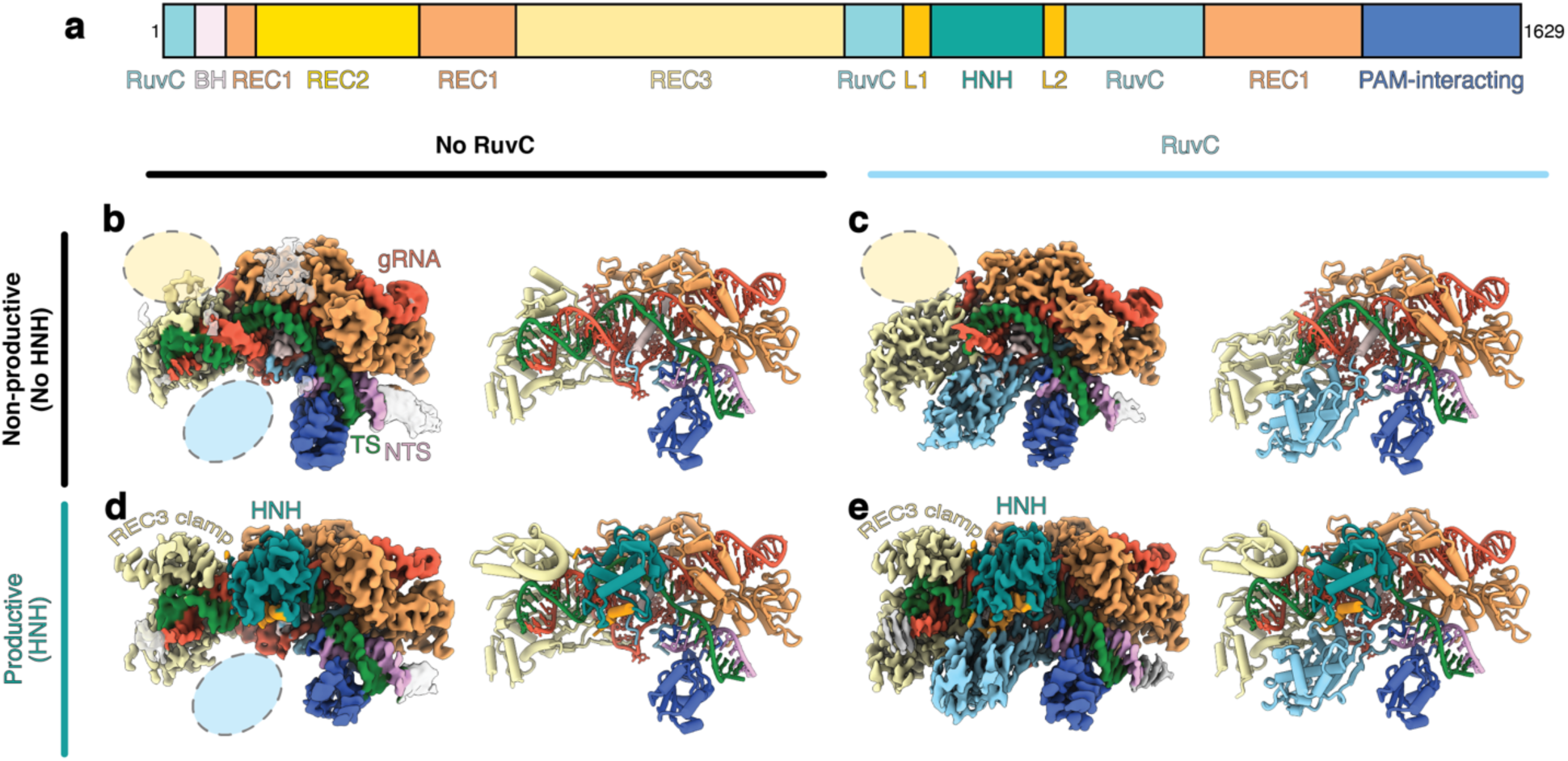
Visualization of distinct stages of FnCas9 nuclease activation. a, FnCas9 domain organization. b, 2.9 Å cryo-EM reconstruction of FnCas9 in a non-productive state with no RuvC density resolved. Yellow oval highlights missing REC3 density, and blue oval highlights missing RuvC density. The corresponding models are depicted to the right of the cryo-EM reconstructions. c, 2.9 Å cryo-EM reconstruction of FnCas9 in a non-productive state with RuvC density. d, 3.0 Å cryo-EM reconstruction of FnCas9 in a productive state with HNH in the active conformation and no RuvC density. e, 2.6 Å cryo-EM reconstruction of FnCas9 in a productive state with RuvC density. The structures are colored as in (a) and as follows: TS, green; NTS, pink; gRNA, red.

**Fig. 2.**
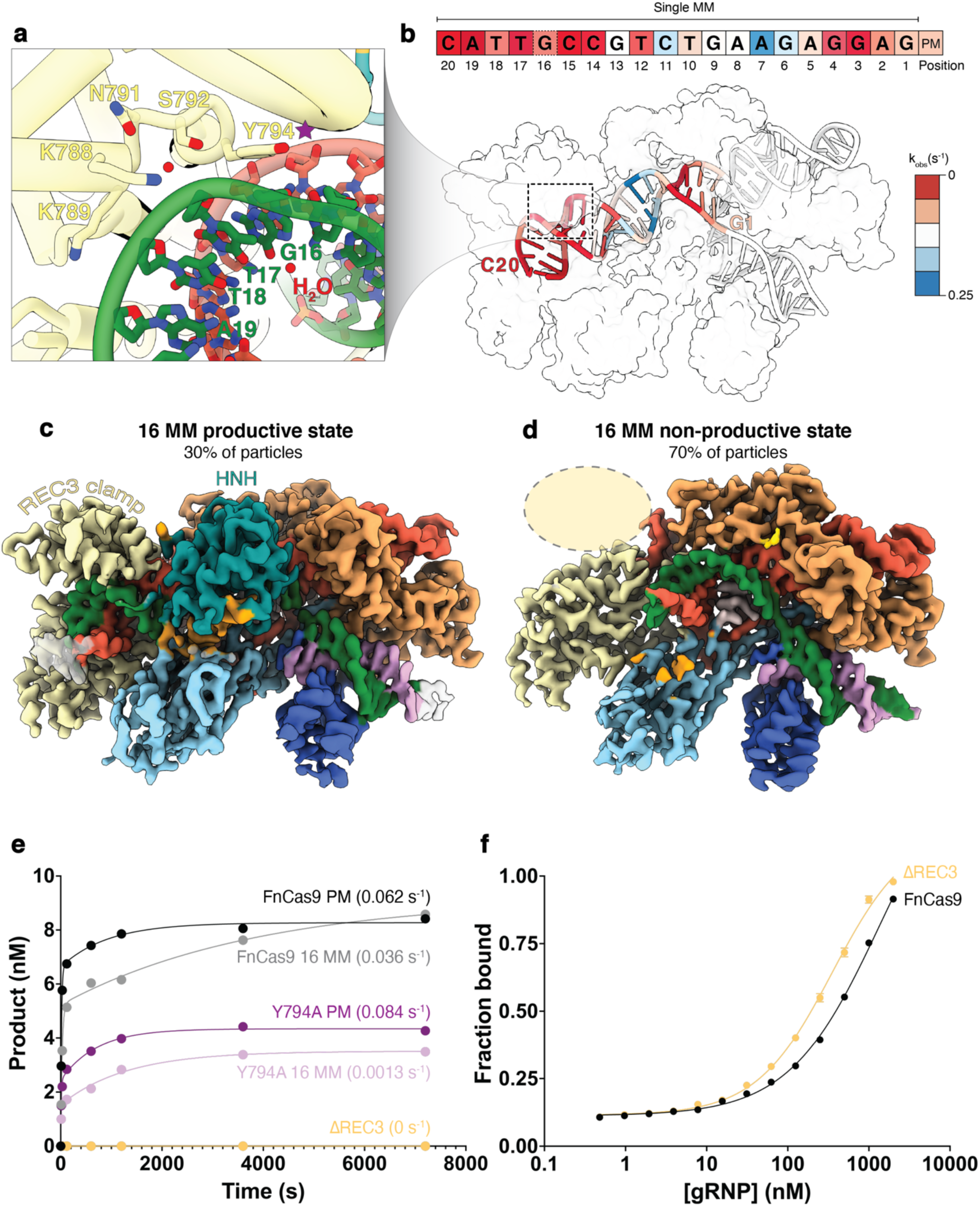
The REC3 clamp enhances FnCas9 specificity. a, Detailed view of REC3 clamp residues contacting the PAM-distal heteroduplex. Purple star indicates Y794 which is subsequently mutated to alanine in e-f. b, Surface representation of FnCas9 in the product state with the nucleotides colored by observed cleavage rate for a single mismatch at that TS position. Observed cleavage rates were measured for each mismatch via capillary electrophoresis. Red indicates a slower observed cleavage rate, and blue indicates a faster observed cleavage rate. c-d, FnCas9 was incubated with DNA containing a mismatch at position 16 of the TS for 1 hour. The reaction was quenched via vitrification and analyzed via cryo-EM. c, 30% of the particles (79,858 particles) from this dataset yielded a 2.9 Å cryo-EM reconstruction of FnCas9 in a productive state bound to DNA with a mismatch at position 16. d, 70% of the particle (128,469 particles) from this dataset comprised a 3.0 Å cryo-EM reconstruction of FnCas9 in a non-productive state bound to DNA with a mismatch at position 16. e, 100 nM of each FnCas9 variant was mixed with 25 nM of each substrate and the cleaved DNA product was monitored via capillary electrophoresis. Observed cleavage rates were calculated using KinTek Explorer. f, Various concentrations of preformed FnCas9:gRNA (RNP) were mixed with 10 nM of on-target DNA and allowed to equilibrate for two hours. Binding was measured by fluorescence polarization.

### Conservation analysis

The on-target FnCas9 structure with RuvC and HNH present was given to ConSurf^30^ for conservation analysis. Homologues were collected from the UniProt^48^ database via a HMMR search^49^. The E-value cutoff was set to 0.001 with three iterations. The CD-HIT cutoff was set to 99%, maximum number of homologues to 150, and minimum sequence identity to 10%. The minimum query sequence coverage was set to 60%. A multiple sequence alignment was then built using MAFFT, and conservation scores calculated by the Bayesian method. The conservation scores were then mapped onto the on-target FnCas9 structure with RuvC and HNH present. The bar graph depicting amino acid position versus conservation score was made using Jalview^50^.

## Results

### FnCas9 undergoes distinct structural changes during nuclease activation

Previous structural studies of FnCas9 were performed using inactive enzyme, where the catalytic residue N995 in the HNH domain was mutated to alanine^22^. Inactivating HNH may hinder structural rearrangements required for enzyme activation^23,24^. To elucidate the conformational changes associated with FnCas9 activation, we prepared cryo-EM grids of fully active FnCas9 in complex with gRNA and 55-bp target DNA after a one-hour incubation. The target DNA substrate contained the preferred PAM for FnCas9, NGG^22^. This single cryo-EM dataset yielded four unique, high-resolution reconstructions of FnCas9 during the process of enzyme activation. (**Fig. 1, Extended Data Fig. 1, Extended Data Fig. 2 and Extended Data Table 1**). The reconstructions were distinguished by the presence or absence of the two catalytic domains, HNH and RuvC.

All four reconstructions demonstrate that FnCas9 recognizes the NGG PAM primarily through hydrogen bonding between G2 and R1585, and G3 and R1556 (**Extended Data Fig. 3**), consistent with previous reports^22^. The two non-productive structures of FnCas9 do not contain resolvable HNH density, indicating that the HNH domain is flexibly tethered and inactive in these states. We also noticed that a large portion of REC3 is missing from the non-productive reconstructions. In the non-productive reconstruction where RuvC is observed, only 13 bp of the R-loop are resolved (**Fig. 1c**). At lower thresholds, the PAM-distal R-loop can be seen sticking out away from the enzyme, hence why it was averaged out in the high-resolution reconstruction (**Extended Data Fig. 4**). In the other non-productive structure without RuvC density, we observe the R-loop in its entirety, including contacts formed between REC3 and the PAM-distal R-loop (**Fig. 1b**). REC3 residue Q801 contacts the gRNA phosphate backbone at position 15 of the R-loop. N791 similarly contacts the TS phosphate backbone, but between R-loop positions 17 and 18. Together, the non-productive structures of FnCas9 point to a checkpoint during enzyme activation where approximately 100 amino acids in REC3 are disordered prior to docking onto the PAM-distal heteroduplex. We call this portion of REC3 the REC3 clamp.

The productive FnCas9 reconstructions from this dataset contain well-resolved HNH domains in the active conformation (**Fig. 1d-e**). Furthermore, the phosphodiester backbone of the TS is cleaved at the anticipated HNH endonuclease site three nucleotides upstream of the PAM. In addition to HNH, the REC3 clamp is well-resolved, and forms stabilizing contacts with the completed R-loop in both productive conformations. In the productive structure lacking RuvC density, we observe flexibility in the portion of REC3 outside of the clamp that typically interfaces with RuvC, although the REC3 clamp remains in contact with the PAM-distal heteroduplex. These findings suggest a stepwise mechanism in which the REC3 clamp must dock onto the fully formed R-loop prior to HNH rearrangement and subsequent DNA cleavage.

### The REC3 clamp confers FnCas9 specificity

SpCas9 is well-documented to tolerate mismatches in the PAM-distal region, leading to cleavage of off-target sequences, which pose a significant challenge in genome editing applications^5,10,25,26^. In contrast, FnCas9 exhibits a markedly higher sensitivity to mismatches in this region, where a single mismatch can abolish cleavage activity^17^. While this intrinsic specificity has been observed, the underlying molecular mechanism has remained elusive. To investigate this, we analyzed the cleavage kinetics of FnCas9 using guide RNA (gRNA) and DNA substrates containing a single mismatch at each position along the protospacer (**Fig. 2b and Extended Data Table 2**).

Our results revealed that mismatches in the PAM-distal region slowed cleavage rates by up to 155-fold compared to perfectly matched substrates (**Fig. 2b and Extended Data Table 3**). Structural inspection of the productive state of FnCas9 uncovered extensive contacts between the REC3 clamp and the PAM-distal heteroduplex, particularly in regions where mismatches significantly impaired cleavage (**Fig. 2a**). These interactions include S792 and K788, which coordinate a water molecule with the target strand (TS) phosphodiester backbone, and K789, which forms electrostatic interactions with the TS backbone at positions 18 and 19. Most notably, Y794 stacks on the ribose at position 16 of the TS, stabilizing the final base pairs of the R-loop (**Fig. 2a and Extended Data Fig. 5**).

To probe the functional relevance of Y794, we performed cryo-EM studies of FnCas9 with gRNA and a DNA substrate with a mismatch at position 16 of the protospacer. These structural studies revealed that only ∼30% of particles adopted a productive state, where the HNH domain had rearranged into the catalytically active conformation, while the remaining ∼70% of particles were trapped in a non-productive state (**Fig. 2c-d**). Despite the mismatch, Y794 maintained its stacking interaction in the productive particles, highlighting the importance of this residue in docking of the REC3 clamp.

We next evaluated the role of Y794 in mismatch discrimination through mutational analysis. Substitution of Y794 with alanine (Y794A) drastically reduced the cleavage rate for off-target substrates with mismatches at position 16, slowing it by 27-fold relative to wild-type FnCas9 (**Fig. 2e**). In addition, the Y794A mutant exhibited a substantial decrease in overall product formation, even at extended time points. These results suggest that Y794-mediated ribose stacking is critical for enabling the REC3 clamp to dock onto the PAM-distal duplex. Without this interaction, approximately half of the Y794A molecules became trapped in a non-productive state, where the REC3 clamp failed to dock and HNH rearrangement could not occur.

Further supporting the essential role of the REC3 clamp, deletion of the REC3 region (Δ450-858) completely abolished cleavage activity, but retained DNA binding (**Fig. 2f**). Together, these findings establish the REC3 clamp as a molecular gate that confers the exceptional specificity of FnCas9. By sensing PAM-distal complementarity, the REC3 clamp imposes an additional checkpoint during nuclease activation, ensuring that only on-target DNA undergoes cleavage.

Unlike SpCas9, which frequently tolerates mismatches in the PAM-distal region, FnCas9 relies on the REC3 clamp to dock onto the final base pairs of the R-loop. This unique mechanism underpins off-target stringency and provides a framework for engineering Cas9 variants with enhanced fidelity.

### Structural basis of heightened virulence in *F. novicida*

Unlike canonical CRISPR-Cas9 systems that target foreign nucleic acid for cleavage, *F. novicida* encodes an additional scaRNA capable of repressing its own genes rather than destroying them^20,27^. In *F. novicida*, scaRNA forms a complex with FnCas9 and tracrRNA to bind an endogenous bacterial lipoprotein (BLP) gene (**Fig. 3a and Extended Data Fig. 7**). Notably, this scaRNA complex does not cleave the BLP gene—an event that would damage its own genome—but instead represses its expression by targeting the 5′UTR. This repression evades host Toll-like receptor 2 detection, reducing immune activation.

**Fig. 3.**
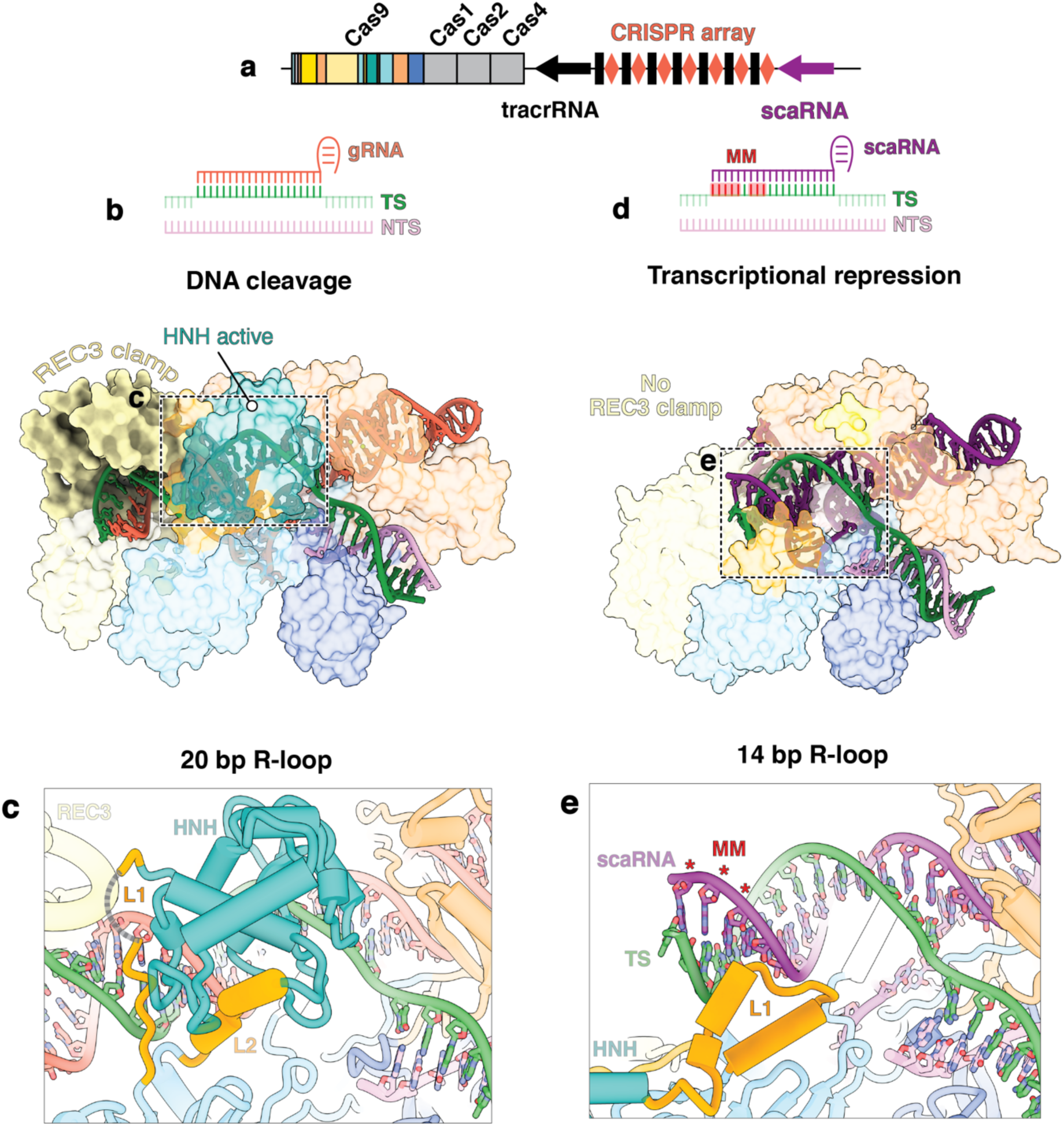
The REC3 clamp senses target complementarity distinguishing DNA cleavage from transcriptional repression. a, Schematic of the *Francisella novicida* chromosomal locus consisting of Cas genes, tracrRNA, crRNA, and scaRNA. b, Schematic of canonical FnCas9 targeting where 20 bp of R-loop form between the gRNA and TS. c, Detailed view of L1 when the R-loop has propagated to completion and HNH is in the active conformation. d, Schematic of scaRNA-mediated targeting where 11 bp of complementarity exists prior to encountering mismatches. 2.8 Å cryo-EM reconstruction of scaRNA-mediated targeting by FnCas9 with 14 bp of R-loop formed. e, Detailed view of L1 in scaRNA-mediated transcriptional repression structure with mismatches designated in red.

To clarify how FnCas9 distinguishes cleavage versus repression of a given target, we solved a cryo-EM structure of FnCas9 bound to a guide RNA that mimics scaRNA and the 5′UTR of *FTN_1101*—one of the genes encoding the BLP (**Fig. 3d and Extended Data Fig. 8**). Our dataset revealed a dominant conformation in which the R-loop extends for 14 base pairs with positions 12–14 mismatched (**Fig. 3e**). This partial R-loop fails to recruit the REC3 clamp, which in turn prevents the HNH domain from adopting a cleavage-competent conformation. We observed the HNH linker 1 (L1) folded in an inactive conformation (**Fig. 3e**), consistent with an R-loop that is distorted and incomplete^10,28^.

In comparison, a fully matched target promotes R-loop completion, permits REC3 clamp docking, and enables HNH domain rearrangement for DNA cleavage (**Fig. 3c**). Thus, our findings reveal that the REC3 clamp imposes an additional activation step, allowing FnCas9 to toggle between cleavage and repression depending on R-loop length. This mechanism underscores how FnCas9 can be repurposed to silence endogenous genes and thereby enhance bacterial virulence without compromising bacterial genome integrity.

### The REC3 clamp is conserved across type II-B CRISPR systems

Type II-B CRISPR-Cas effectors are known to be larger than other type II enzymes^29^, prompting us to question whether these systems share the extended REC3 domain observed in FnCas9, and possibly a similar mismatch-sensing mechanism. To address this question, we performed a ConSurf conservation analysis of 51 type II-B Cas9 homologs^30^, revealing that the key REC3 clamp residues are as highly conserved as the catalytic residues in the HNH and RuvC domains (**Fig. 4a–c**). Notably, Q798, which intercalates between the guide RNA and TS, S792, which coordinates a water molecule at the TS backbone, and Y794, which stacks on the target-strand ribose at position 16, all show strong sequence conservation.

**Fig. 4.**
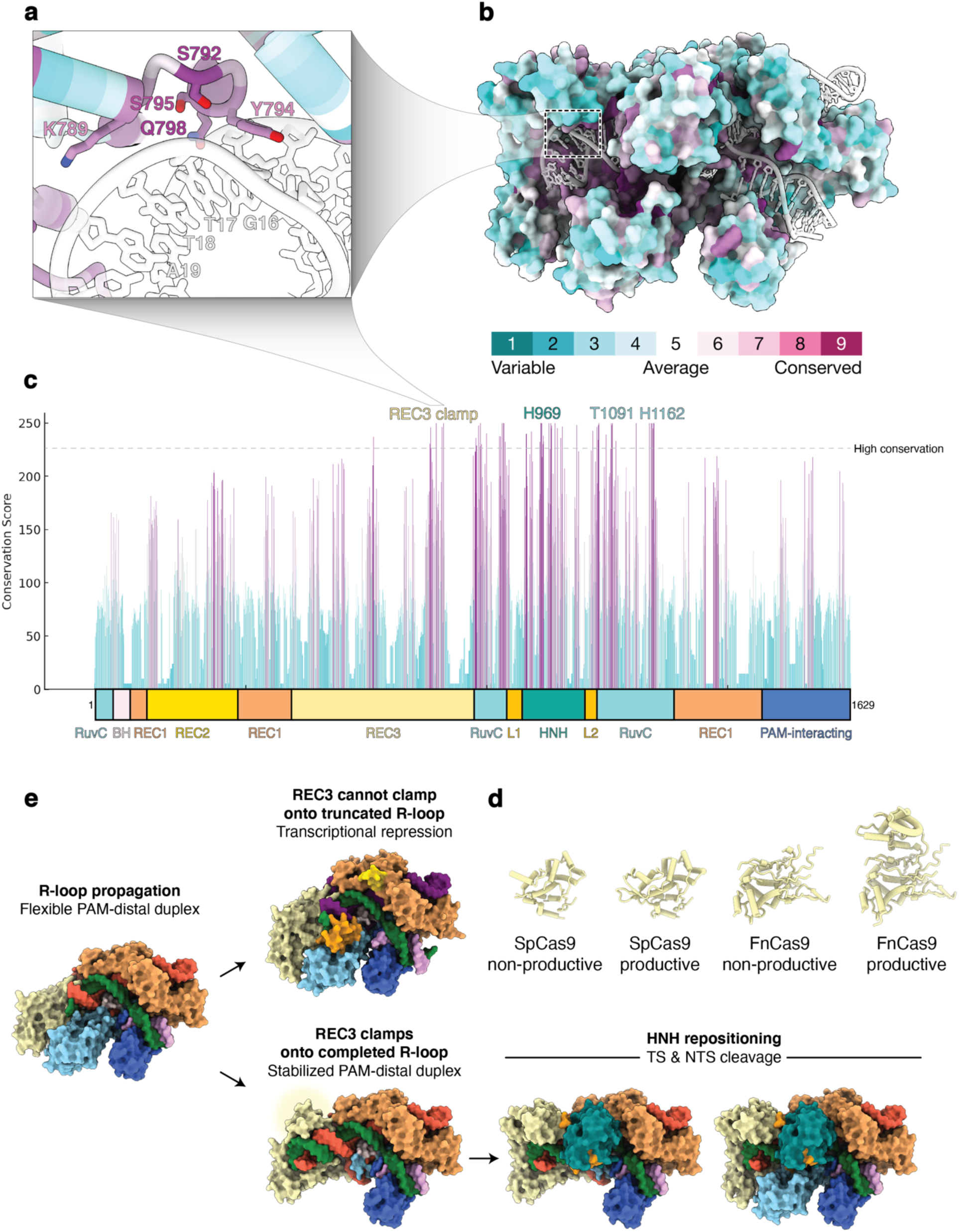
Type II-B CRISPR-Cas systems contain extended REC3 domains that add an additional checkpoint during enzyme activation. a, Detailed view of REC3 clamp residues contacting the PAM-distal heteroduplex colored by conservation score. The location of these residues is highlighted in b-c. b, Surface representation of FnCas9 in the product state where each residue is colored by conservation, where purple represents more conserved residues, and cyan denotes more variable residues. c, Domain schematic of FnCas9 with the corresponding conservation score for each residue. Residues with a conservation score of 225 or higher are considered highly conserved. d, Comparison of REC3 domains from SpCas9 in non-productive (PDB 7S4U) and productive states (PDB 7S4X)^10^, and FnCas9 in non-productive and productive states. e, Model of type II-B CRISPR-Cas9 nuclease activation. If the R-loop propagates to completion, the REC3 clamp docks onto the PAM-distal heteroduplex prompting HNH repositioning and DNA cleavage. Alternatively, if REC3 cannot clamp due to incomplete R-loop formation, type II-B CRISPR-Cas9 repress transcription without cleavage.

In contrast, SpCas9 (a type II-A enzyme) contains a smaller REC3 domain (**Fig. 4d and Extended Data Fig. 9**), which lacks the extensive contacts seen in FnCas9 for sensing the final base pairs of the R-loop. This difference correlates with the established tolerance of SpCas9 for PAM-distal mismatches^5,10,25,26^, whereas the extended REC3 clamp of FnCas9 imposes an additional checkpoint that strictly hinders off-target cleavage. Our findings suggest that type II-B Cas9 enzymes contain extended REC3 domains capable of sensing PAM-distal mismatches, thereby imposing an additional activation step that minimizes off-target cleavage (**Fig. 4e**).

In *F. novicida*, the REC3 clamp also underlies the ability of the enzyme to repress rather than cleave specific endogenous genes, thus facilitating immune evasion through subverted host detection. Altogether, the REC3 clamp emerges as a hallmark of type II-B Cas9 systems, simultaneously driving heightened specificity and virulence. Its unique properties distinguish type II-B enzymes from other CRISPR-Cas systems and offer a promising framework for designing next-generation genome editors with improved fidelity and tunable outcomes.

## Discussion

Our work uncovers a novel specificity checkpoint in FnCas9, centered on the REC3 clamp, that transforms our understanding of how type II-B CRISPR-Cas9 orthologs achieve heightened fidelity. Distinct cryo-EM reconstructions reveal a stepwise activation process, where the R-loop must propagate to completion before the REC3 clamp docks, enabling HNH domain rearrangement, and DNA cleavage. In contrast, SpCas9 is known to efficiently cleave DNA with mismatches, particularly in the PAM-distal region. This discrepancy arises because SpCas9 lacks a REC3 clamp, enabling L1 to dock onto the heteroduplex once the R-loop has propagated to ∼16 bp, thereby activating the HNH domain despite mismatches. This also underlies the rapid rate of SpCas9 DNA cleavage. However, in FnCas9 the REC3 clamp remains undocked if mismatches are present, preventing L1 from engaging the heteroduplex and halting HNH activation. This clamp mechanism explains why FnCas9 is so intolerant to PAM-distal mismatches—a common blind spot to SpCas9 (**Extended Data Fig. 10**)^10,31^.

Beyond specificity, we also show how the REC3 clamp differentiates transcriptional repression from cleavage, evidenced by the scaRNA complex that targets the *F. novicida* chromosome. The REC3 clamp remains undocked when the R-loop is incomplete, leaving HNH in a non-productive conformation. From a pathogenesis standpoint, this mechanism allows *F. novicida* to avoid host immune recognition by silencing critical BLP genes without risking self-inflicted genome damage.

The clamp and its high conservation among type II-B CRISPR-Cas9 enzymes have practical implications. First, the clamp domain could be engineered into other Cas9 variants or smaller Cas effectors to minimize off-target cleavage^32–35^. Second, the reliance of *F. novicida* and related pathogens on REC3 clamp-based repression suggests new antibacterial strategies, such as specifically targeting the clamp or scaRNA function to disrupt bacterial immune evasion and sensitize pathogens to immune clearance or antibiotics ^21,36–38^.

Overall, our findings provide a blueprint for advanced genome editing tools that harness the REC3 clamp for improved precision. They also highlight a previously underappreciated strategy in bacterial virulence that might be exploited to counter antibiotic resistance. As new CRISPR-based applications emerge, understanding the modular features that influence both cleavage and repression will be key to expanding our therapeutic and biotechnological toolkit.

## Acknowledgements

We thank J.P.K. Bravo and the Taylor group for insightful discussions. We are also grateful to the members of the group of K.A. Johnson.

## Author Contributions

Conceptualization: G.N.H. Investigation and data analysis: G.N.H. (protein production, cryo-EM data collection, structure determination and modeling, biochemistry, kinetic analysis, AF3 structural prediction, and conservation analysis). Writing initial draft: G.N.H. G.N.H. and D.W.T. reviewed, edited, and approved the manuscript. Supervision: D.W.T. Funding acquisition: D.W.T.

## Data Availability

The structures and associated atomic coordinates have been deposited into the Electron Microscopy Data Bank (EMDB) and Protein Data Bank (PDB) with accession codes FnCas9 productive (EMD-48052 and PDB 9EHF), FnCas9 productive no RuvC (EMD-48053 and PDB 9EHG), FnCas9 non-productive (EMD-48054 and PDB 9EHH), FnCas9 non-productive no RuvC (EMD-48062 and PDB 9EHR), FnCas9 16 MM productive (EMD-48070 and PDB 9EHX), FnCas9 16 MM non-productive (EMD-48069 and PDB 9EHW), and FnCas9 scaRNA gRNA 1101 DNA non-productive (EMD-49074 and PDB 9N6T).

## References

1. Sorek, R., Lawrence, C. M. & Wiedenheft, B. CRISPR-mediated adaptive immune systems in bacteria and archaea. Annu Rev Biochem 82, 237–266 (2013).

2. Jinek, M. et al. A programmable dual-RNA-guided DNA endonuclease in adaptive bacterial immunity. Science (1979) 337, 816–821 (2012).

3. Mali, P. et al. RNA-guided human genome engineering via Cas9. Science (1979) 339, 823–826 (2013).

4. Cong, L. et al. Multiplex genome engineering using CRISPR/Cas systems. Science (1979) 339, 819–823 (2013).

5. Fu, Y. et al. High-frequency off-target mutagenesis induced by CRISPR-Cas nucleases in human cells. Nature Biotechnology 2013 31:9 31, 822–826 (2013).

6. Doudna, J. A. The promise and challenge of therapeutic genome editing. Nature 2020 578:7794 578, 229–236 (2020).

7. Kleinstiver, B. P. et al. High-fidelity CRISPR–Cas9 nucleases with no detectable genome-wide off-target effects. Nature 2015 529:7587 529, 490–495 (2016).

8. Chen, J. S. et al. Enhanced proofreading governs CRISPR–Cas9 targeting accuracy. Nature 2017 550:7676 550, 407–410 (2017).

9. Slaymaker, I. M. et al. Rationally engineered Cas9 nucleases with improved specificity. Science (1979) 351, 84–88 (2016).

10. Bravo, J. P. K. et al. Structural basis for mismatch surveillance by CRISPR– Cas9. Nature 2022 603:7900 603, 343–347 (2022).

11. Slaymaker, I. M. & Gaudelli, N. M. Engineering Cas9 for human genome editing. Curr Opin Struct Biol 69, 86–98 (2021).

12. Russa, M. F. La & Qi, L. S. The New State of the Art: Cas9 for Gene Activation and Repression. Mol Cell Biol 35, 3800–3809 (2015).

13. Thakore, P. I. et al. RNA-guided transcriptional silencing in vivo with S. aureus CRISPR-Cas9 repressors. Nature Communications 2018 9:1 9, 1–9 (2018).

14. Wang, J. et al. Engineering a PAM-flexible SpdCas9 variant as a universal gene repressor. Nature Communications 2021 12:1 12, 1–10 (2021).

15. Konermann, S. et al. Genome-scale transcriptional activation by an engineered CRISPR-Cas9 complex. Nature 2014 517:7536 517, 583–588 (2014).

16. McCarty, N. S., Graham, A. E., Studená, L. & Ledesma-Amaro, R. Multiplexed CRISPR technologies for gene editing and transcriptional regulation. Nature Communications 2020 11:1 11, 1–13 (2020).

17. Acharya, S. et al. Francisella novicida Cas9 interrogates genomic DNA with very high specificity and can be used for mammalian genome editing. Proc Natl Acad Sci U S A 116, 20959–20968 (2019).

18. Sternberg, S. H., Lafrance, B., Kaplan, M. & Doudna, J. A. Conformational control of DNA target cleavage by CRISPR–Cas9. Nature 2015 527:7576 527, 110–113 (2015).

19. Singh, D. et al. Mechanisms of improved specificity of engineered Cas9s revealed by single-molecule FRET analysis. Nature Structural & Molecular Biology 2018 25:4 25, 347–354 (2018).

20. Sampson, T. R., Saroj, S. D., Llewellyn, A. C., Tzeng, Y. L. & Weiss, D. S. A CRISPR-CAS System Mediates Bacterial Innate Immune Evasion and Virulence. Nature 497, 254 (2013).

21. Ratner, H. K. et al. Catalytically Active Cas9 Mediates Transcriptional Interference to Facilitate Bacterial Virulence. Mol Cell 75, 498–510.e5 (2019).

22. Hirano, H. et al. Structure and Engineering of Francisella novicida Cas9. Cell 164, 950–961 (2016).

23. Sun, W. et al. Structures of Neisseria meningitidis Cas9 Complexes in Catalytically Poised and Anti-CRISPR-Inhibited States. Mol Cell 76, 938 (2019).

24. Pacesa, M. et al. R-loop formation and conformational activation mechanisms of Cas9. Nature 2022 609:7925 609, 191–196 (2022).

25. Hsu, P. D. et al. DNA targeting specificity of RNA-guided Cas9 nucleases. Nature Biotechnology 2013 31:9 31, 827–832 (2013).

26. Kuscu, C., Arslan, S., Singh, R., Thorpe, J. & Adli, M. Genome-wide analysis reveals characteristics of off-target sites bound by the Cas9 endonuclease. Nature Biotechnology 2014 32:7 32, 677–683 (2014).

27. Ratner, H. K. et al. Catalytically Active Cas9 Mediates Transcriptional Interference to Facilitate Bacterial Virulence. Mol Cell 75, 498–510.e5 (2019).

28. Zhang, Y. et al. Catalytic-state structure and engineering of Streptococcus thermophilus Cas9. Nature Catalysis 2020 3:10 3, 813–823 (2020).

29. Chylinski, K., Makarova, K. S., Charpentier, E. & Koonin, E. V. Classification and evolution of type II CRISPR-Cas systems. Nucleic Acids Res 42, 6091 (2014).

30. Yariv, B. et al. Using evolutionary data to make sense of macromolecules with a “face-lifted” ConSurf. Protein Science 32, e4582 (2023).

31. Pacesa, M. et al. Structural basis for Cas9 off-target activity. Cell 185, 4067–4081.e21 (2022).

32. Harrington, L. B. et al. Programmed DNA destruction by miniature CRISPR-Cas14 enzymes. Science (1979) 362, 839–842 (2018).

33. Ma, D. et al. Engineer chimeric Cas9 to expand PAM recognition based on evolutionary information. Nature Communications 2019 10:1 10, 1–9 (2019).

34. Xu, X. et al. Engineered miniature CRISPR-Cas system for mammalian genome regulation and editing. Mol Cell 81, 4333–4345.e4 (2021).

35. Zhao, L. et al. PAM-flexible genome editing with an engineered chimeric Cas9. Nature Communications 2023 14:1 14, 1–8 (2023).

36. Sampson, T. R., Saroj, S. D., Llewellyn, A. C., Tzeng, Y. L. & Weiss, D. S. A CRISPR/Cas system mediates bacterial innate immune evasion and virulence. Nature 2013 497:7448 497, 254–257 (2013).

37. Djordjevic, M. et al. In silico analysis suggests common appearance of scaRNAs in type II systems and their association with bacterial virulence. Front Genet 9, 17 (2018).

38. Xu, P. X. et al. Distribution characteristics of the Legionella CRISPR-Cas system and its regulatory mechanism underpinning phenotypic function. Infect Immun 92, (2024).

39. Hibshman, G. N. et al. Unraveling the mechanisms of PAMless DNA interrogation by SpRY-Cas9. Nature Communications 2024 15:1 15, 1–15 (2024).

40. Dangerfield, T. L., Huang, N. Z. & Johnson, K. A. High throughput quantification of short nucleic acid samples by capillary electrophoresis with automated data processing. Anal Biochem 629, 114239 (2021).

41. Mastronarde, D. N. Automated electron microscope tomography using robust prediction of specimen movements. J Struct Biol 152, 36–51 (2005).

42. Punjani, A. Real-time cryo-EM structure determination. Microscopy and Microanalysis 27, 1156–1157 (2021).

43. Jumper, J. et al. Highly accurate protein structure prediction with AlphaFold. Nature 2021 596:7873 596, 583–589 (2021).

44. Emsley, P., Lohkamp, B., Scott, W. G. & Cowtan, K. Features and development of Coot. Acta Crystallogr D Biol Crystallogr 66, 486 (2010).

45. Croll, T. I. ISOLDE: A physically realistic environment for model building into low-resolution electron-density maps. Acta Crystallogr D Struct Biol 74, 519–530 (2018).

46. Adams, P. D. et al. PHENIX: a comprehensive Python-based system for macromolecular structure solution. Acta Crystallogr D Biol Crystallogr 66, 213 (2010).

47. Pettersen, E. F. et al. UCSF ChimeraX: Structure visualization for researchers, educators, and developers. Protein Science 30, 70–82 (2021).

48. Bateman, A. et al. UniProt: the universal protein knowledgebase. Nucleic Acids Res 45, D158–D169 (2017).

49. Finn, R. D., Clements, J. & Eddy, S. R. HMMER web server: interactive sequence similarity searching. Nucleic Acids Res 39, W29 (2011).

50. Waterhouse, A. M., Procter, J. B., Martin, D. M. A., Clamp, M. & Barton, G. J. Jalview Version 2-A multiple sequence alignment editor and analysis workbench. Bioinformatics 25, 1189–1191 (2009).

