## Extended Data for "Structural basis for a dual-function type II-B CRISPR-Cas9"

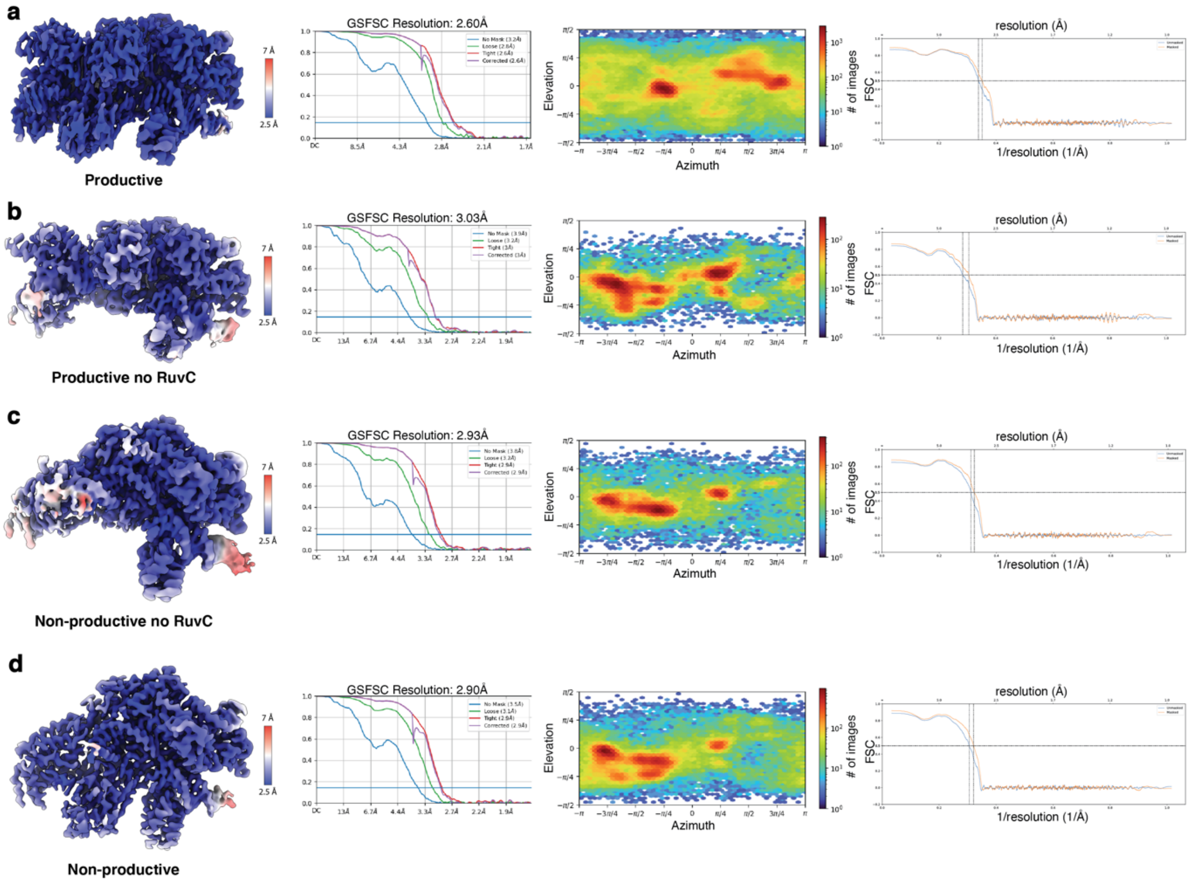


**Extended Data Figure 1:** Cryo-EM data analysis of FnCas9 bound to PM DNA in the a, productive b, productive no RuvC c, non-productive no RuvC and d, non-productive states.

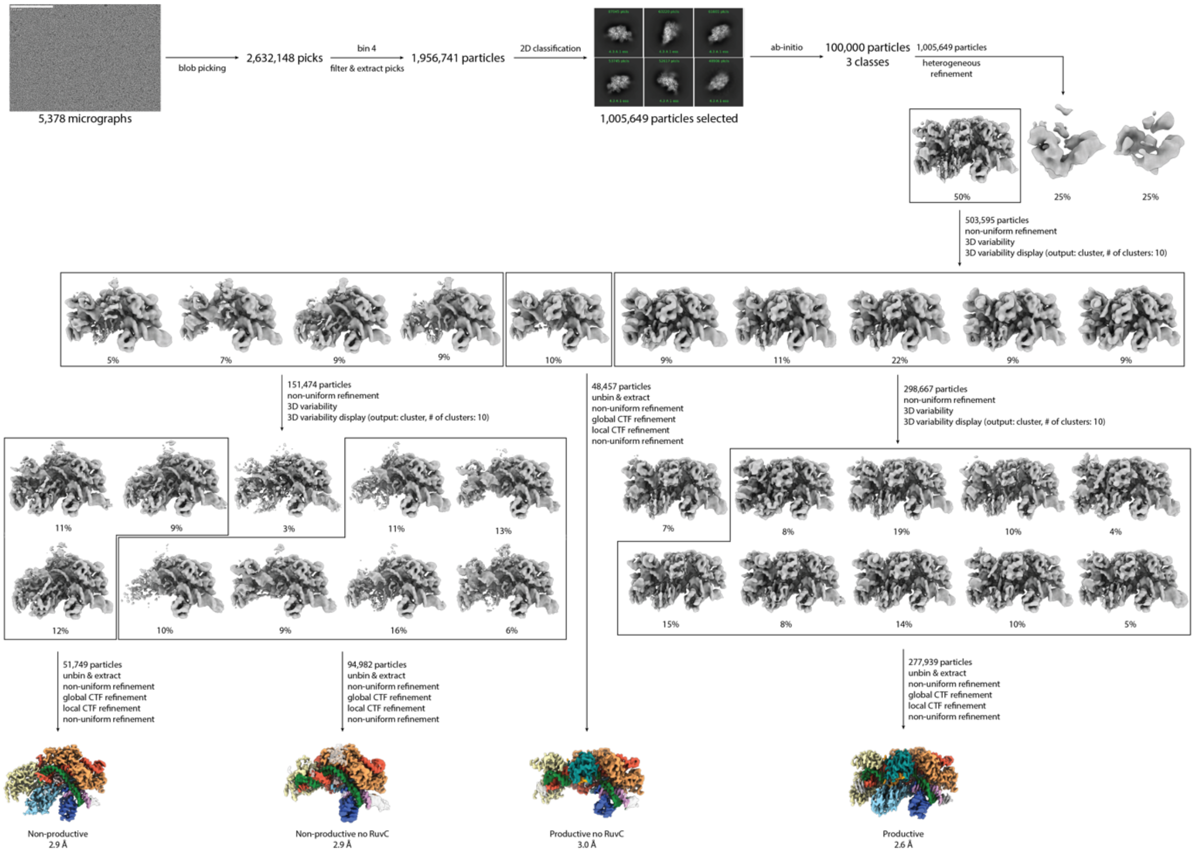


**Extended Data Figure 2:** Cryo-EM data processing workflow for FnCas9 bound to PM DNA after a 1-hour incubation at room temperature. Final reconstructions are colored by domain.

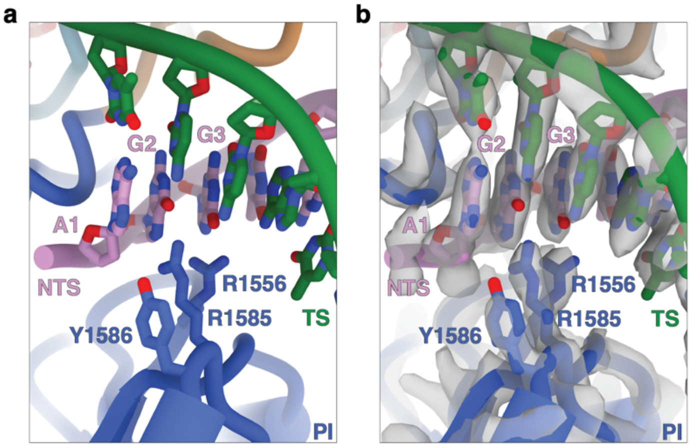


**Extended Data Figure 3:** FnCas9 residues R1556 and R1585 recognize the NGG PAM. a, Detailed view of the PAM site interaction when FnCas9 is in the product state. b, Cryo-EM density of the FnCas9 PAM site interaction.

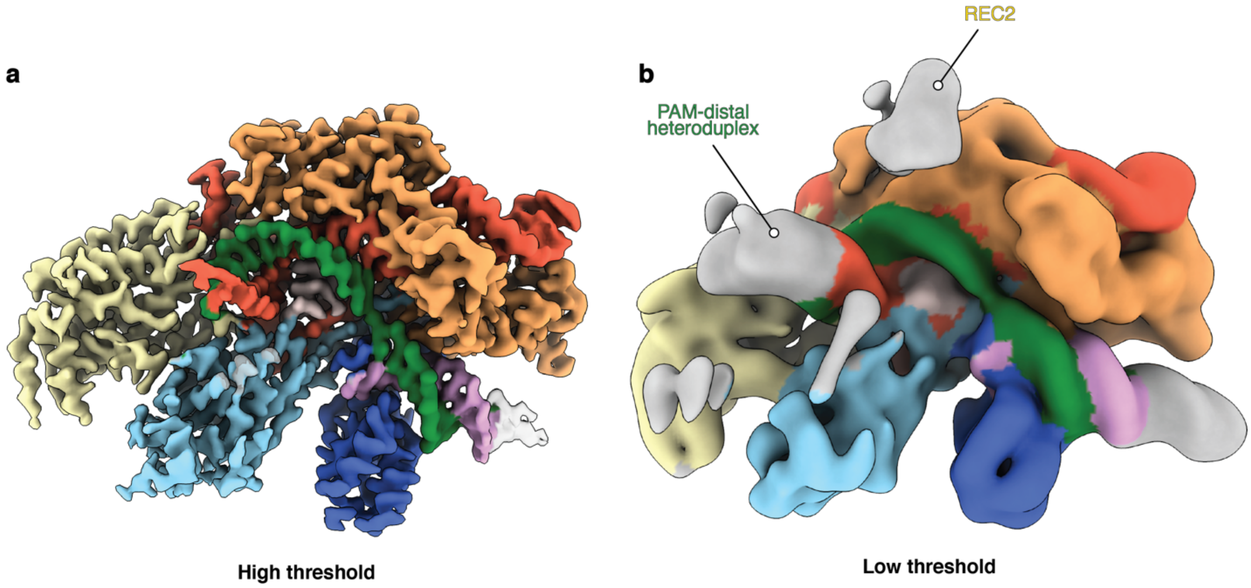


**Extended Data Figure 4:** The PAM-distal heteroduplex is flexible prior to REC3 docking. a, 2.9 Å cryo-EM reconstruction of FnCas9 in a non-productive conformation at a high threshold where the PAM-distal heteroduplex is averaged out, and b, at a low threshold where the PAM-distal heteroduplex is resolved.

a b

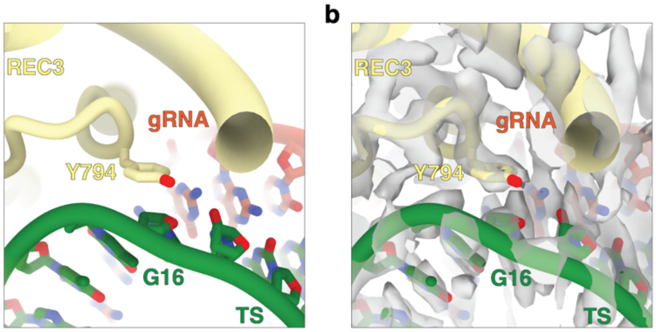


**Extended Data Figure 5:** REC3 clamp residue Y794 stacks upon the ribose at position 16 of the TS. a, Detailed view of the REC3 ribose stacking interaction when FnCas9 is in the product state. b, Cryo-EM density for the REC3 ribose stacking interaction at 2.6 Å resolution.

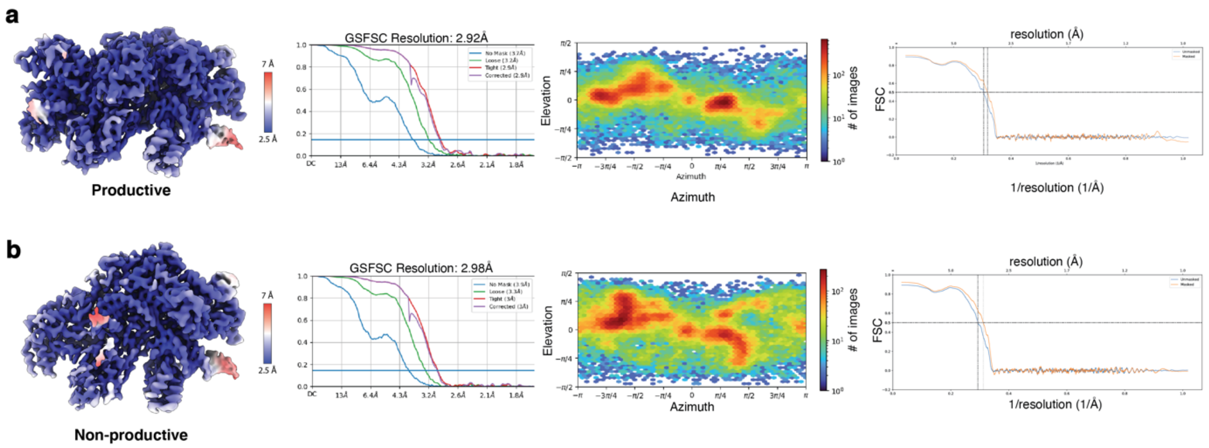


**Extended Data Figure 6:** Cryo-EM data analysis of FnCas9 bound to 16 MM DNA in the a, productive and b, non-productive states.

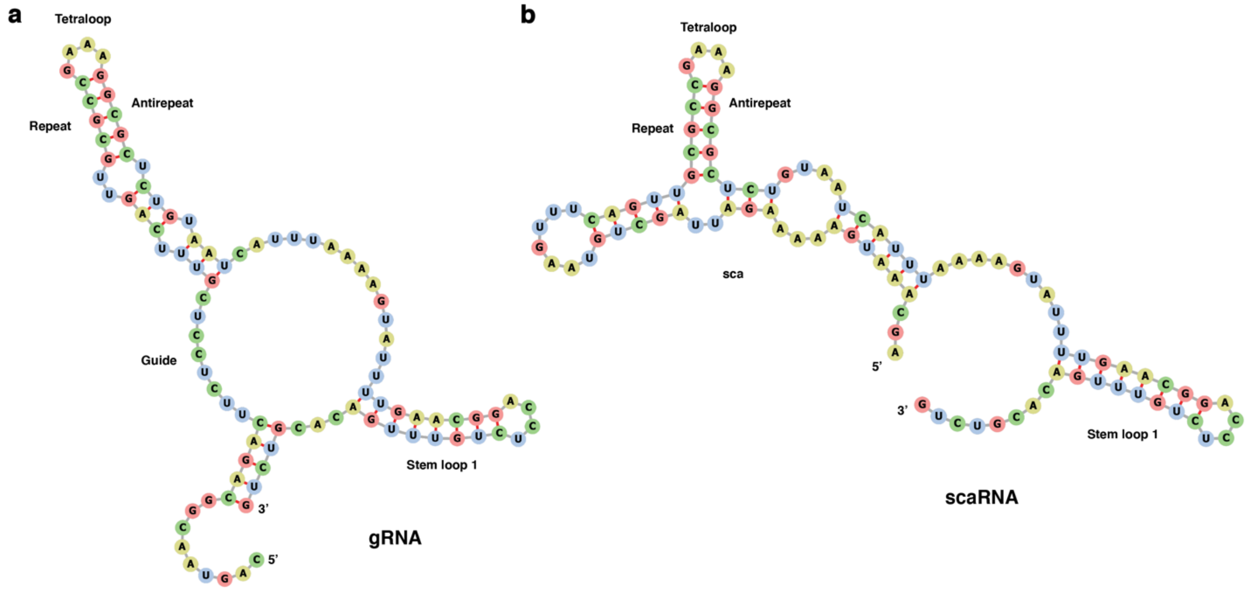


**Extended Data Figure 7:** Comparison between FnCas9 a, gRNA and b, scaRNA single guide mimic secondary structures.

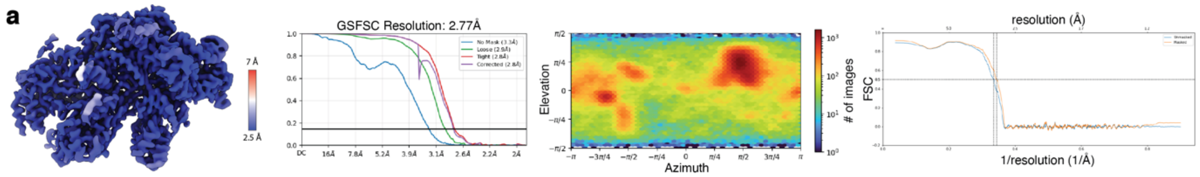


**Extended Data Figure 8:** Cryo-EM data analysis of FnCas9 bound to scaRNA gRNA targeting 1101 DNA in the a, non-productive state.

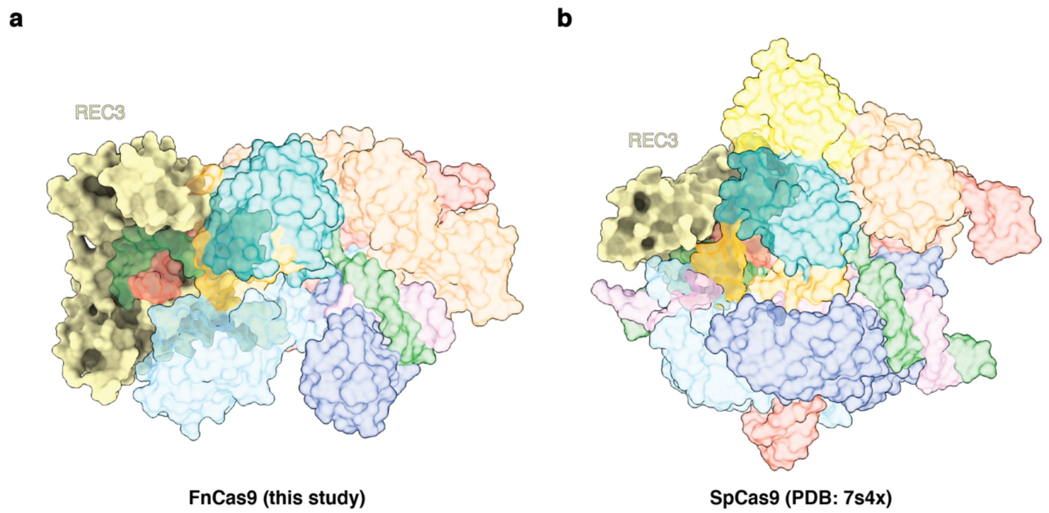


**Extended Data Figure 9:** FnCas9 and SpCas9 REC3 domain comparison. a, FnCas9 residues 450-858 comprise the REC3 domain. b, SpCas9 residues 496-718 comprise the REC3 domain. REC3 domains are opaque.

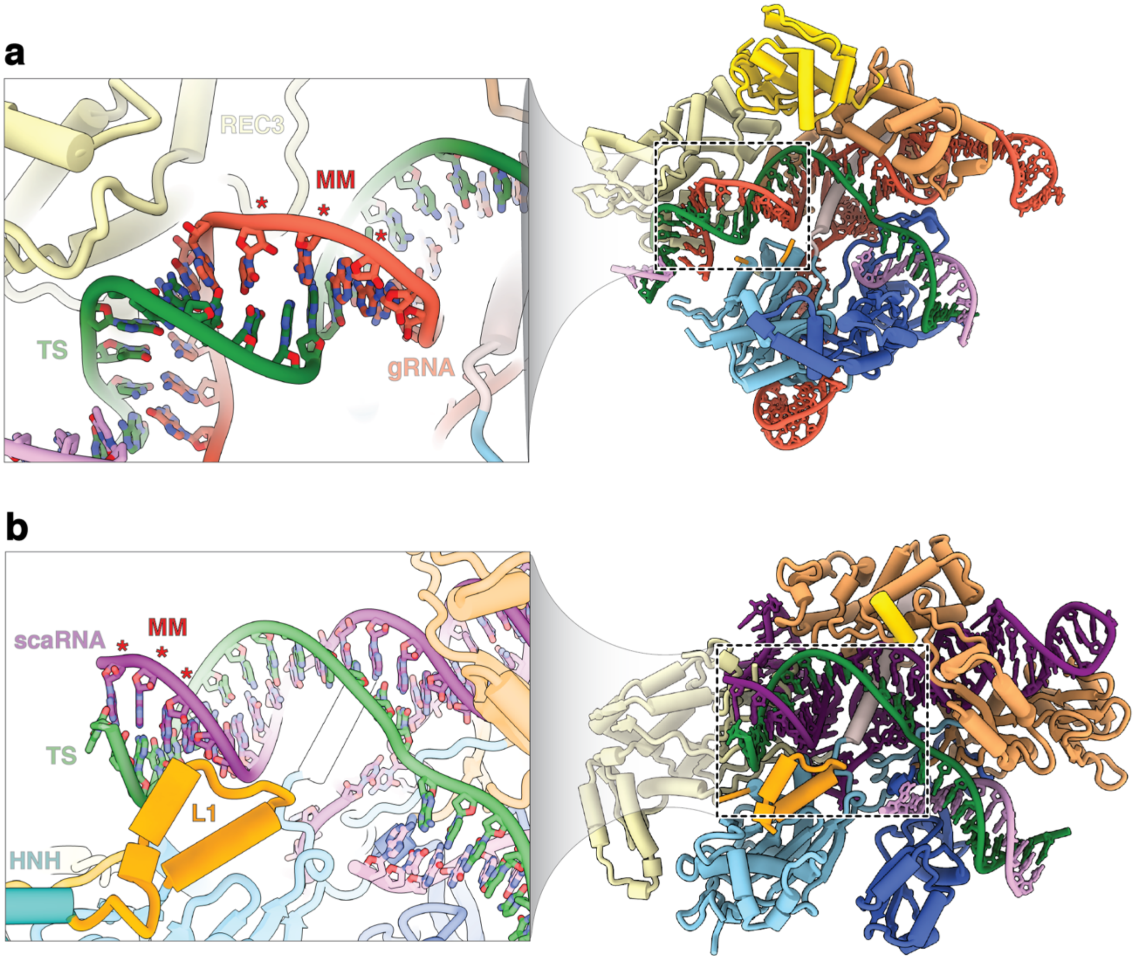


**Extended Data Figure 10:** Comparison of SpCas9 and FnCas9 12-14 MM surveillance structures. a, SpCas9 bound to 12-14 MM DNA highlighting the lack of REC3 residues contacting this region of the heteroduplex. REC3 is able to dock onto this mismatched DNA. b, FnCas9 scaRNA-mediated transcriptional repression structure with positions 12-14 mismatched. REC3 is unable to dock onto the truncated heteroduplex.

| | FnCas9 PM Productive RuvC (EMD-48052) (PDB 9EHF) | FnCas9 PM Productive No RuvC (EMD-48053) (PDB 9EHG) | FnCas9 PM Non-Productive RuvC (EMD-48054) (PDB 9EHH) | FnCas9 PM Non-Productive No RuvC (EMD-48062) (PDB 9EHR) | FnCas9 16MM Productive (EMD-48070) (PDB 9EHX) | FnCas9 16MM Non-Productive (EMD-48069) (PDB 9EHW) | FnCas9 scaRNA gRNA 1101 DNA Non-productive (EMD-49074) (PDB 9N6T) |
|---|---|---|---|---|---|---|---|
| **Data collection and processing** | | | | | | | |
| Voltage (kV) | 300 kV | 300 kV | 300 kV | 300 kV | 300 kV | 300 kV | 200 kV |
| Electron exposure (e–/Å$^2$) | 80 | 80 | 80 | 80 | 80 | 80 | 50 |
| Defocus range (μm) | -1.5 to -2.5 | -1.5 to -2.5 | -1.5 to -2.5 | -1.5 to -2.5 | -1.5 to -2.5 | -1.5 to -2.5 | -1.5 to -2.5 |
| Pixel size (Å) | 0.8332 | 0.8332 | 0.8332 | 0.8332 | 0.8332 | 0.8332 | 0.933 |
| Symmetry imposed | C1 | C1 | C1 | C1 | C1 | C1 | C1 |
| Initial particle images (no.) | 1,956,741 | 1,956,741 | 1,956,741 | 1,956,741 | 2,495,949 | 2,495,949 | 1,573,500 |
| Final particle images (no.) | 277,939 | 48,457 | 51,749 | 94,982 | 79,858 | 63,288 | 283,943 |
| Map resolution (Å) | 2.6 | 3.0 | 2.9 | 2.9 | 2.9 | 3.0 | 2.9 |
| FSC threshold | 0.5 | 0.5 | 0.5 | 0.5 | 0.5 | 0.5 | 0.5 |
| Map resolution range (Å) | 2.5-7 | 2.5-7 | 2.5-7 | 2.5-7 | 2.5-7 | 2.5-7 | 2.5-7 |
| **Refinement** | | | | | | | |
| Initial model used (PDB code) | 5B2O | 9EHF | 9EHF | 9EHG | 9EHF | 9EHH | AF3 model |
| Model resolution (Å) | 2.8 | 3.0 | 3.1 | 3.1 | 3.1 | 3.2 | 2.9 |
| | 0.5 | 0.5 | 0.5 | 0.5 | 0.5 | 0.5 | 0.5 |
| FSC threshold | | | | | | | |
| Model resolution range (Å) | | | | | | | |
| Map sharpening *B* factor (Å$^2$) | 70.2 | 67.6 | 72.5 | 80.1 | 79.3 | 78.2 | 88.3 |
| Model composition | | | | | | | |
| Non-hydrogen | 13,919 | 12,107 | 11,230 | 10,211 | 13,917 | 11,158 | 12,009 |
| atoms | 1,356 | 1,123 | 1,069 | 891 | 1,356 | 1,057 | 1,134 |
| Protein residues | 133 | 133 | 116 | 133 | 133 | 117 | 125 |
| Nucleotides | 2 $Mg^{2+}$ | 2 $Mg^{2+}$ | 2 $Mg^{2+}$ | 2 $Mg^{2+}$ | 2 $Mg^{2+}$ | 2 $Mg^{2+}$ | 0 |
| Ligands | | | | | | | |
| R.m.s. deviations | | | | | | | |
| Bond lengths (Å) | 0.005 | 0.006 | 0.005 | 0.006 | 0.005 | 0.005 | 0.005 |
| Bond angles (°) | 0.943 | 1.153 | 1.034 | 1.000 | 1.023 | 1.094 | 0.970 |
| Validation | | | | | | | |
| MolProbity score | 0.99 | 1.01 | 0.96 | 1.02 | 0.95 | 0.93 | 0.83 |
| Clashscore | 1.81 | 2.27 | 1.96 | 2.36 | 1.77 | 1.73 | 1.16 |
| Poor rotamers (%) | 0.99 | 0.98 | 0.42 | 0.98 | 0.25 | 0.84 | 0.29 |
| Ramachandran plot | | | | | | | |
| Favored (%) | 97.77 | 98.29 | 98.10 | 98.86 | 97.92 | 98.47 | 98.04 |
| Allowed (%) | 2.23 | 1.71 | 1.90 | 1.14 | 2.08 | 1.53 | 1.96 |
| Disallowed (%) | 0.00 | 0.00 | 0.00 | 0.00 | 0.00 | 0.00 | 0.00 |

**Extended Data Table 1:** Cryo-EM data collection, refinement and validation statistics.

| Name | Sequence (5’-3’) | Source |
|---|---|---|
| HBB_TS | /6-FAM/CATGGTGCATCTGACTCCTGAGGAGAAGTCTGCCGTTACTGCCCTGTGGGGCAAG | IDT |
| HBB_NTS | CTTGCCCCACAGGGCAGTAACGGCAGACTTCTCCTCAGGAGTCAGATGCACCATG | IDT |
| HBB_TS_1 | /6-FAM/CATGGTGCATCTGACTCCTGAGGAGAAGTCTGCCGTTAGTGCCCTGTGGGGCAAG | IDT |
| HBB_NTS_1 | CTTGCCCCACAGGGCACTAACGGCAGACTTCTCCTCAGGAGTCAGATGCACCATG | IDT |
| HBB_TS_2 | /6-FAM/CATGGTGCATCTGACTCCTGAGGAGAAGTCTGCCGTTTCTGCCCTGTGGGGCAAG | IDT |
| HBB_NTS_2 | CTTGCCCCACAGGGCAGAAACGGCAGACTTCTCCTCAGGAGTCAGATGCACCATG | IDT |
| HBB_TS_3 | /6-FAM/CATGGTGCATCTGACTCCTGAGGAGAAGTCTGCCGTAACTGCCCTGTGGGGCAAG | IDT |
| HBB_NTS_3 | CTTGCCCCACAGGGCAGTTACGGCAGACTTCTCCTCAGGAGTCAGATGCACCATG | IDT |
| HBB_TS_4 | /6-FAM/CATGGTGCATCTGACTCCTGAGGAGAAGTCTGCCGATACTGCCCTGTGGGGCAAG | IDT |
| HBB_NTS_4 | CTTGCCCCACAGGGCAGTATCGGCAGACTTCTCCTCAGGAGTCAGATGCACCATG | IDT |
| HBB_TS_5 | /6-FAM/CATGGTGCATCTGACTCCTGAGGAGAAGTCTGCCCTTACTGCCCTGTGGGGCAAG | IDT |
| HBB_NTS_5 | CTTGCCCCACAGGGCAGTAAGGGCAGACTTCTCCTCAGGAGTCAGATGCACCATG | IDT |
| HBB_TS_6 | /6-FAM/CATGGTGCATCTGACTCCTGAGGAGAAGTCTGCGGTTACTGCCCTGTGGGGCAAG | IDT |
| HBB_NTS_6 | CTTGCCCCACAGGGCAGTAACCGCAGACTTCTCCTCAGGAGTCAGATGCACCATG | IDT |
| HBB_TS_7 | /6-FAM/CATGGTGCATCTGACTCCTGAGGAGAAGTCTGGCGTTACTGCCCTGTGGGGCAAG | IDT |
| HBB_NTS_7 | CTTGCCCCACAGGGCAGTAACGCCAGACTTCTCCTCAGGAGTCAGATGCACCATG | IDT |
| HBB_TS_8 | /6-FAM/CATGGTGCATCTGACTCCTGAGGAGAAGTCTCCCGTTACTGCCCTGTGGGGCAAG | IDT |
| HBB_NTS_8 | CTTGCCCCACAGGGCAGTAACGGGAGACTTCTCCTCAGGAGTCAGATGCACCATG | IDT |
| HBB_TS_9 | /6-FAM/CATGGTGCATCTGACTCCTGAGGAGAAGTCAGCCGTTACTGCCCTGTGGGGCAAG | IDT |
| HBB_NTS_9 | CTTGCCCCACAGGGCAGTAACGGCTGACTTCTCCTCAGGAGTCAGATGCACCATG | IDT |
| HBB_TS_10 | /6-FAM/CATGGTGCATCTGACTCCTGAGGAGAAGTGTGCCGTTACTGCCCTGTGGGGCAAG | IDT |
| HBB_NTS_10 | CTTGCCCCACAGGGCAGTAACGGCACACTTCTCCTCAGGAGTCAGATGCACCATG | IDT |
| HBB_TS_11 | /6-FAM/CATGGTGCATCTGACTCCTGAGGAGAAGACTGCCGTTACTGCCCTGTGGGGCAAG | IDT |
| HBB_NTS_11 | CTTGCCCCACAGGGCAGTAACGGCAGTCTTCTCCTCAGGAGTCAGATGCACCATG | IDT |
| HBB_TS_12 | /6-FAM/CATGGTGCATCTGACTCCTGAGGAGAACTCTGCCGTTACTGCCCTGTGGGGCAAG | IDT |
| HBB_NTS_12 | CTTGCCCCACAGGGCAGTAACGGCAGAGTTCTCCTCAGGAGTCAGATGCACCATG | IDT |
| HBB_TS_13 | /6-FAM/CATGGTGCATCTGACTCCTGAGGAGATGTCTGCCGTTACTGCCCTGTGGGGCAAG | IDT |
| HBB_NTS_13 | CTTGCCCCACAGGGCAGTAACGGCAGACATCTCCTCAGGAGTCAGATGCACCATG | IDT |
| HBB_TS_14 | /6-FAM/CATGGTGCATCTGACTCCTGAGGAGTAGTCTGCCGTTACTGCCCTGTGGGGCAAG | IDT |

| | | |
|---|---|---|
| HBB_NTS_14 | CTTGCCCCACAGGGCAGTAACGGCAGACTACTCCTCAGGAGTCAGATGCACCATG | IDT |
| HBB_TS_15 | /6-FAM/CATGGTGCATCTGACTCCTGAGGACAAGTCTGCCGTTACTGCCCTGTGGGGCAAG | IDT |
| HBB_NTS_15 | CTTGCCCCACAGGGCAGTAACGGCAGACTTGTCCTCAGGAGTCAGATGCACCATG | IDT |
| HBB_TS_16 | /6-FAM/CATGGTGCATCTGACTCCTGAGGTGAAGTCTGCCGTTACTGCCCTGTGGGGCAAG | IDT |
| HBB_NTS_16 | CTTGCCCCACAGGGCAGTAACGGCAGACTTCACCTCAGGAGTCAGATGCACCATG | IDT |
| HBB_TS_17 | /6-FAM/CATGGTGCATCTGACTCCTGAGCAGAAGTCTGCCGTTACTGCCCTGTGGGGCAAG | IDT |
| HBB_NTS_17 | CTTGCCCCACAGGGCAGTAACGGCAGACTTCTGCTCAGGAGTCAGATGCACCATG | IDT |
| HBB_TS_18 | /6-FAM/CATGGTGCATCTGACTCCTGACGAGAAGTCTGCCGTTACTGCCCTGTGGGGCAAG | IDT |
| HBB_NTS_18 | CTTGCCCCACAGGGCAGTAACGGCAGACTTCTCGTCAGGAGTCAGATGCACCATG | IDT |
| HBB_TS_19 | /6-FAM/CATGGTGCATCTGACTCCTGTGGAGAAGTCTGCCGTTACTGCCCTGTGGGGCAAG | IDT |
| HBB_NTS_19 | CTTGCCCCACAGGGCAGTAACGGCAGACTTCTCCACAGGAGTCAGATGCACCATG | IDT |
| HBB_TS_20 | /6-FAM/CATGGTGCATCTGACTCCTCAGGAGAAGTCTGCCGTTACTGCCCTGTGGGGCAAG | IDT |
| HBB_NTS_20 | CTTGCCCCACAGGGCAGTAACGGCAGACTTCTCCTGAGGAGTCAGATGCACCATG | IDT |
| HBB_TS_PT | /6-FAM/CATGGTGCATCTGACTCCTGAGGAGAAGTCTGCC*G*T*T*A*CTGCCCTGTGGGGCAAG | IDT |
| HBB_NTS_PT | CTTGCCCCACAGGGCAG*T*A*A*C*GGCAGACTTCTCCTCAGGAGTCAGATGCACCATG | IDT |
| 1101_TS | ATTATCATCCTTATTTTTTTATTATTATTAACATTGGATTTCTCCATTACAGCTAATTACTAAGTAAATAGC | IDT |
| 1101_NTS | /6-FAM/GCTATTTACTTAGTAATTAGCTGTAATGGAGAAATCCAATGTTAATAATAATAAAAAAATAAGGATGATAAT | IDT |
| scaRNA_gRNA | AGUAGGCGACGUAUAAUGUGGUUUCAGUUGCGCCGAAAGGCGCUCUGUAAUCAUUUAAAAGUAUUUUGAACGGACCUCUGUUUGACACGUCUG | IDT |
| HBB_gRNA | CAGUAACGGCAGACUUCUCCUCGUUUCAGUUGCGCCGAAAGGCGCUCUGUAAUCAUUUAAAAGUAUUUUGAACGGACCUCUGUUUGACACGUCUG | IDT |

**Extended Data Table 2:** List of nucleotide sequences used in this study.

| **DNA substrate** | **$k_{obs}$ ($s^{-1}$)** |
|---|---|
| PM | 0.066 |
| 1 MM | 0.074 |
| 2MM | 0.045 |
| 3 MM | 0.004 |
| 4 MM | 0.007 |
| 5 MM | 0.078 |
| 6 MM | 0.149 |
| 7 MM | 0.222 |
| 8 MM | 0.102 |
| 9 MM | 0.099 |
| 10 MM | 0.086 |
| 11 MM | 0.148 |
| 12 MM | 0.027 |
| 13 MM | 0.101 |
| 14 MM | 0.012 |
| 15 MM | 0.001 |
| 16 MM | 0.036 |
| 17 MM | 0.001 |
| 18 MM | 0.034 |
| 19 MM | 0.001 |
| 20 MM | 0.001 |

**Extended Data Table 3:** FnCas9 observed cleavage rates for perfect match, and single mismatch (MM) DNA substrates.
